# Death comes for us all: an interplay of habitat selection, movement, and social behavior relate to cause specific mortality among grey wolves

**DOI:** 10.1101/2021.03.02.433605

**Authors:** Julie W. Turner, Christina M. Prokopenko, Katrien A. Kingdon, Daniel L. J. Dupont, Sana Zabihi-Seissan, Eric Vander Wal

**Affiliations:** Memorial University of Newfoundland, Department of Biology, 232 Elizabeth Ave., St. John’s, NL Canada A1B 3X9; Manitoba, Wildlife and Fisheries Branch, 200 Saulteaux Cresc., Winnipeg, MB Canada R3J 3W3

**Keywords:** *Canis lupus*, anthropogenic disturbance, disease, integrated step selection analysis, animal behavior, canine distemper virus

## Abstract

Avoiding death infects biological processes, including behavior. Habitat selection, movement, and sociality are highly flexible behaviors that influence the mortality risks and subsequent fitness of individuals. In the Anthropocene, animals are experiencing increased risks from direct human causes and increased spread of infectious diseases. Using integrated step selection analysis, we tested how the habitat selection, movement, and social behaviors of grey wolves vary as an individual dies due to humans or canine distemper virus (CDV) and how those behaviors may vary in the lead up to death. Behaviors that changed prior to death were strongly related to how an animal eventually died. Wolves killed by humans moved slower than wolves that survived and selected to be nearer roads closer in time to their death. Wolves that died due to CDV moved progressively slower as they neared death and reduced their avoidance of wet habitats. All animals, regardless of dying or not maintained strong selection to be near packmates across time, which seemingly contributed to disease dynamics in the packs that became infected with CDV. Habitat selection, movement, and sociality interact to put individuals and groups at greater risks, influencing their cause-specific mortality.

**Lay Summary:** Not much is known about behaviors prior to death in wild animals. Grey wolves killed by humans selected to be in riskier areas increasingly prior to their deaths. Wolves that died due to disease moved slower and changed their habitat selection to be in areas with more water as they became sicker. Sick wolves also continued to select for packmates, increasing the chances that the whole pack would succumb to the disease.

## Introduction

Death comes for us all and fundamentally shapes behavior as animals attempt to survive in environments composed of risks. Mortality drives demography, and to mitigate this risk animals modulate their behaviors. Fine-scale behaviors, such as habitat selection, movement, and sociality, are flexible in many animal species, and animals have evolved to adjust their behavioral responses to minimize long-term and short-term risks. Some of the most common causes of death to large, highly mobile animals are humans – ranging from hunting to vehicle collisions – and disease (Farias et al. 2005; M.H. Murray and St Clair 2015; Wynn-Grant et al. 2018). Anthropogenic risks have become inescapable as the wildlife-human interface expands (Sih et al. 2010; Skelhorn et al. 2011; Haswell et al. 2017). Many predators alter their behaviors to avoid the long-term impacts of anthropogenic disturbance by minimizing human-wildlife conflict both spatially and temporally (Broekhuis et al. 2019; Nickel et al. 2019). With increasing human disturbance, the risk of disease outbreaks also grows, which may induce behavioral change, or sickness behaviors (Weary et al. 2009; Becker et al. 2020). Being ill creates short-term risks that lead to temporary changes in behavior. Symptomatic animals often move less, adjust their space use, and change their social patterns, particularly in more gregarious species (Hart 2011). An individual’s mortality is intertwined with habitat selection, movement, and social behaviors (Webber and Vander Wal 2017), and here we test how these behaviors relate to the specific cause of mortality in a social carnivore.

Habitat selection, where animals preferentially use different habitats in a non-random manner, is one way animals can alter their behaviors that can have demographic consequences (Morris 2003). Animals change their habitat selection based on the density of conspecifics, dynamic resource availability, and predation risks (McLoughlin et al. 2010; Avgar et al. 2020). Selection of complex habitats by predators is associated with their higher mortality due to prey availability or predation risk (Gigliotti et al. 2020). Many species adjust their habitat selection to avoid human disturbance, particularly during the day when humans are most active (Martin et al. 2010; Gaynor et al. 2019 Feb 8). Humans frequently kill predators when they are near their livestock or trapped/hunted as fur-bearer species, which are typically in open areas or along trails (Recio et al. 2018). Sick animals also trade-off gaining easier access to food by selecting riskier habitats that have human provisioning (M. Murray et al. 2015; M.H. Murray and St Clair 2017).

Movement behavior is necessary for variation in space use as habitat selection is possible because of an individual’s ability to travel between habitat types (Forester et al. 2007). Movement is mechanistically determined by the internal states, motions, and navigation capabilities of individuals (Nathan et al. 2008), all of which are dynamic and can be altered depending on risks (Knowlton and Graham 2010). For example, mule deer (*Odocoileus hemionus*) that migrated along central, high-use routes had much greater survival than those that traveled more peripheral routes (Sawyer et al. 2019). Individuals migrating along the periphery were depredated more often than individuals travelling more centrally or in bigger groups (Sawyer et al. 2019). Moreover, many species tend to move faster in anthropogenic areas to avoid risks from humans (Tucker et al. 2018). Diseased or injured banded mongooses (*Mungos mungo*) are less active, move less, and delay dispersal (Fairbanks, Hawley, and Alexander 2014a; Fairbanks, Hawley, and Alexander 2014b). Successfully delaying dispersal when recovering from injury or illness can enables the individual to avoid death and maintain reproductive opportunities that may have been lost if they did not alter their behavior (Boydston et al. 2005; Altizer et al. 2011). Dispersal, for example, demonstrates how animals cannot preferentially use different areas without movement and results in fitness consequences ranging from obtaining enough nutrients to survive to reproductive opportunities to mitigating more direct mortality risks.

Variations in sociality can also have important fitness ramifications (J.B. Silk 2007). It is well established in many primate species that being more social generally improves fitness, but there is growing evidence for longevity being negatively influenced by sociality in certain situations (reviewed in Thompson 2019 Mar 21). Being social can also counteract negative effects of stress (Kikusui et al. 2006), predation and fluctuating environments (Guindre-Parker and Rubenstein 2020), yet social behavior can be costly in terms of competition and disease risk (Eberhard1975 1975). In response to this sociality trade-off, individuals can behave in a manner that improves their individual fitness, as in healthy house finches (*Carpodacus mexicanus*) actively avoiding sick individuals (Zylberberg et al. 2012). In addition, individuals can exhibit behaviors that improve inclusive fitness through acting in the interest of the group (Cantor et al. 2020). For example, several species of insects demonstrated self-imposed social isolation from the colony when they were near death to mitigate disease risks to the colony (Heinze and Walter 2010; Geffre et al. 2020). However, in social mammals, social isolation is more subtle to observe because the alternations to social interactions occur at a fine scale, such as decreasing physical contact. These adjustments confer the same benefits to their social groups, by maintaining or improving inclusive fitness (Buck et al. 2018; Shakhar 2019). In social species in particular, social behavior is linked to habitat selection and movement behavior as individuals navigate both their social and spatial environments and associated risks (Webber and Vander Wal 2017; He et al. 2019).

Wolves (*Canis lupus*) are a highly mobile social species that are exposed to increasing mortality risks (D.L. Murray et al. 2010; Sidorovich et al. 2017; Joly et al. 2019; Hebblewhite and Whittington 2020). Wolf habitat selection, movement, and social behaviors are highly flexible both throughout the year and in response to rapid environmental change and disturbances (Houle et al. 2010; Mancinelli et al. 2019). Humans frequently kill predators in general, and wolves in particular, when they occur near livestock or as fur-bearer species, typically in open areas or along trails (Theuerkauf 2009). There is evidence that wolves respond to this risk by moving faster in open areas and near human development (Recio et al. 2018). Furthermore, as wolves increasingly use anthropogenic areas, they are at greater risk of contracting diseases such as canine distemper virus (CDV) that frequently occur as reservoirs in urban-rural transition zones (Stronen, Sallows, et al. 2011; Prager et al. 2012; Beineke et al. 2015).

Here, we tested how movement, habitat selection, and sociality related to mortality sources and change as individuals near death using data from GPS collared grey wolves. In the months preceding death, we tested if movement, habitat selection, and sociality would vary predictably, such that the individual’s cause of death would be apparent. Specifically, we tested how movement, habitat selection, and sociality vary prior to death from two types of mortality sources: 1) humans – a long-term risk in the life of an individual and 2) disease – a relatively short-term set of risks while an individual is symptomatic. Specifically, we hypothesized that intrinsic behavioral qualities of individuals, consistent habitat selection, movement, or social behavior patterns, put them at greater risk from humans compared to those that survived (Table S1). Furthermore, we hypothesized that individuals change their habitat selection, movement, and sociality behaviors due the extrinsic influences of disease (Table S1). Thus, we aim to address how variation in habitat selection, movement, and sociality influences not only how an individual dies but also have the potential to change as an animal nears the end of life.

## Methods

### Study sites and subjects

We studied two populations of wolves in in Manitoba, Canada: Riding Mountain National Park (RMNP) and a provincial management unit, Game Hunting Area 26 (GHA 26). Riding Mountain National Park (RMNP; 50°51′50″N 100°02′10″W) is a 2,969 km^2^ protected area in southwestern Manitoba. RMNP is located at the confluence of prairie grassland, aspen parkland and boreal transition, creates a distinct edge with surrounding agricultural land. The RMNP wolf population was estimated from aerial and snow track surveys at ~70 individuals 2016-2017. GHA 26 is an ~7,200 km^2^ unit located in southeastern Manitoba, bordered by Lake Winnipeg to the west and the Manitoba-Ontario border to the east. The landscape is composed predominantly of coniferous and mixed forests, interspersed with rock outcrops, rivers, lakes and bogs. The area is predominantly public land, but also encompasses several communities as well as the Nopiming and Manigotagan River Provincial Parks. The GHA 26 wolf population was estimated to be ~140 wolves by aerial survey between 2014 and 2016.

### Wolf GPS collar data collection

Between 2014 and 2018 wolves in both study areas were captured and fit with GPS telemetry collars (Lotek Iridium TrackM 2D, Lotek Wireless Inc, Newmarket, ON, Canada; Sirtrack Pinnacle G5C, Sirtrack Limited, Hawkes Bay, New Zealand; Followit Tellus Medium 2D, Followit Sweden AB, Lindesberg, Sweden). In RMNP, a total of 25 wolves were captured and collared in winter 2016 and 2017, and 38 wolves in GHA 26 in the winters of 2014-2018. All captures followed Memorial University AUP 16-02-EV. GPS collar data was rarified to a 2-hour relocation schedule to sample animals at an equal intensity for analyses.

In RMNP, 17 of our collared individuals had known causes of death: 40% disease (CDV), 16% anthropogenic (trapping, poisoning, gunshot), and 12% conspecifics (injuries from canids). In GHA 26, 10 of the collared individuals had known causes of death: 18% anthropogenic, 5% disease (heartworm), and 3% drowning. If cause of death was not easily determined in the field (i.e., bullet wound), the cause of death was determined by necropsy. Necropsies were performed by the Assiniboine Park Zoo and Canadian Wildlife Health Cooperative. We categorized death into two main overarching sources of mortality: humans and CDV. As a control, we used 16 wolves that did not die and assigned them a random “death” date that fell in the range of our observed death dates and compared their habitat selection, movement, and social patterns in the time leading up to that date to wolves with verified death dates. Henceforth, we call wolves that succumb to these mortality factors 1) control, 2) human-killed, or 3) CDV wolves.

Canine distemper virus (CDV) is a disease common in carnivores that is very contagious, mainly through direct contact with infected fluids (Ferry 1911). Untreated diseased animals frequently lose motor control and become fevered and dehydrated within one month from infection, so many animals seek out water and may eventually die submersed in water (Blixenkrone-Møller et al. 1993; Rodeheffer et al. 2007; Zhao et al. 2015). Individuals usually die within 2-4 weeks of infection (Loots et al. 2017). Thus, we limited our analyses to the two months leading up to the death date such that control, human-killed, and CDV wolves had at least 1 month of asymptomatic behaviors so we could examine changes over time.

### Integrated step selection analysis

We used integrated step selection analysis (iSSA) to test how habitat selection, movement, and sociality may relate to death, and how these behaviors change at fine temporal scales as an animal nears death. Movement processes are described by two covariates indicating speed and directionality: 1) step length, the straight-line distance between two consecutive GPS locations, and 2) turn angle, the angular deviation from the previous step to the next (Fortin et al. 2005; Avgar et al. 2016). Here, random steps were drawn from theoretical distributions, using gamma for step length (mean tentative shape = 0.398, scale = 2410) and von Mises for turn angle (mean tentative kappa = 0.0957). To ascertain how cause of death related to these behaviors, each variable was interacted with mortality source: control (n = 16), human (n = 4), or CDV (n = 8). To examine how behaviors change leading up to death, we interacted variables of interest with time to death (days). We used natural logs of time to death to standardize it with other variables. Thus, covariates interacted with time to death represented changes in behaviors, and the non-interacted variables represented their intrinsic baseline behaviors.

Tracks, random steps, and covariates for each step were extracted using the ‘amt’ package (Signer et al. 2019) in R v. 3.6.2 (R Core Team 2019). We fit our iSSA model using the ‘glmmTMB’ package (Brooks et al. 2017) using the mixed effects model approach with a Poisson error distribution described in Muff et al. 2019 (). To ensure model convergence, we only included individuals that had >1% availability of a covariate during this two-month period, which has previously been shown to provide a robust representation of the habitat available to individuals (Dickie et al. 2019). Thus, 28 wolves met our criteria for inclusion (RMNP = 14 wolves, GHA 26 = 14 wolves). To ascertain the coefficients of specific individuals in addition to the population, we further included random effects for the slope of each variable for each individual (Muff et al. 2019). When there are a large number of GPS relocations, as we have over our two month observation windows, as few as two animals are sufficient to obtain robust habitat selection estimates (Street et al. 2021).

Our model included variables important to wolf space-use and pertinent to testing our hypotheses. To account for general space-use behaviors known of wolves, we included natural log (ln) of step length (modifier of the gamma shape parameter) alone and interacted the ln step length with proportion of habitat type to account for how animals move differently (faster or slower) through different habitat types. In addition, we included the cosine (cos) of turn angle (von Mises concentration parameter) to describe an individual’s deviation in directionality.

To address our hypotheses, we included covariates to represent 1) movement, 2) habitat selection, and 3) social behaviors in the model. We expected human-killed wolves to have greater speed and selection for human infrastructure as they neared death, but there is no reason to expect them to demonstrate different selection for proximity to packmates compared to control wolves (Theuerkauf 2009). We expected that wolves dying from CDV would decrease movement speed, increase selection for wet habitats, and increase avoidance of packmates as they neared the end of life.

Habitat selection was described by habitat type and distance from road. We defined habitat using the 2015 Canada landcover map at a 30 m × 30 m scale, reclassified to forest, open, and wet habitat types (Table S2). We then calculated the proportion of each habitat type within a 100 m buffer around GPS locations. We included ln distance to roads as an indicator of human presence in the area. Lastly, we included distances to nearest neighbor from the same pack and distance from pack boundary as social covariates. All distance variables were ln transformed because we expect animals to have a stronger response to features when they are closer to them, and that response would decay at an unknown rate.

We calculated nearest neighbor with a collar and the distance to that individual using the ‘spatsoc’ package in R (Robitaille et al. 2019 May 24). For each available point, we calculated the distance to the same neighbor that was the observed nearest neighbor. Because the collared wolves made many forays outside of pack boundaries, we conservatively determined pack boundaries based on wolf behaviors determined by tracking and site investigation where we determined that the wolves hunted, scavenged, or rested when the collars were deployed. We employed kernel density estimations using the center points of these behavioral clusters to approximate pack boundaries using the adehabitatHR package in R (Calenge 2006). Furthermore, we ran a binomial regression to ascertain if they spent more time outside of their pack boundary dependent on their eventual cause of death.

### Relative selection strength

To determine the effect size of selection, we calculated relative selection strength (RSS) of each selection covariate, i.e., forest, open, wet, distance to road, distance to nearest neighbor, and distance to pack boundary (Avgar et al. 2017). Following the method presented in ‘amt’ to calculate the logRSS, we compared the average habitats used to the range of habitats used, one variable at a time (Signer et al. 2019). Specifically, we held habitat variables constant while time to death varied from 61 to 0 days from death.

## Results

General patterns emerged in the population’s habitat selection behavior, wolves moved more in open areas (Table 1) and showed higher relative selection strength (logRSS) for proximity to their nearest packmate than they did habitat variables (Figs. 1 and 2). However, wolves demonstrated a wide range of individual variation in selecting for habitat types and the nearest distances to packmates (Figs. 1 and 2).

**Table 1.**
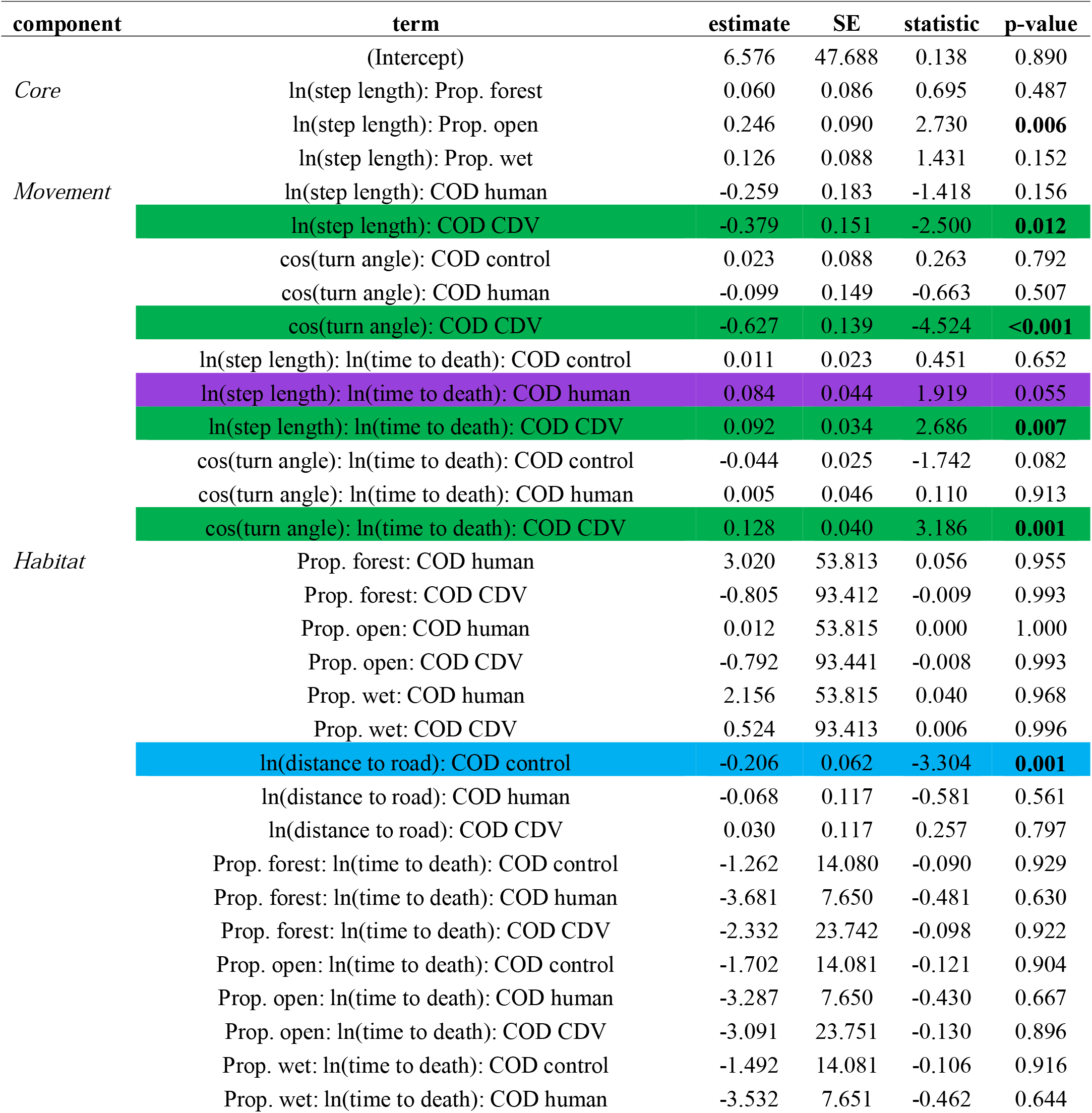
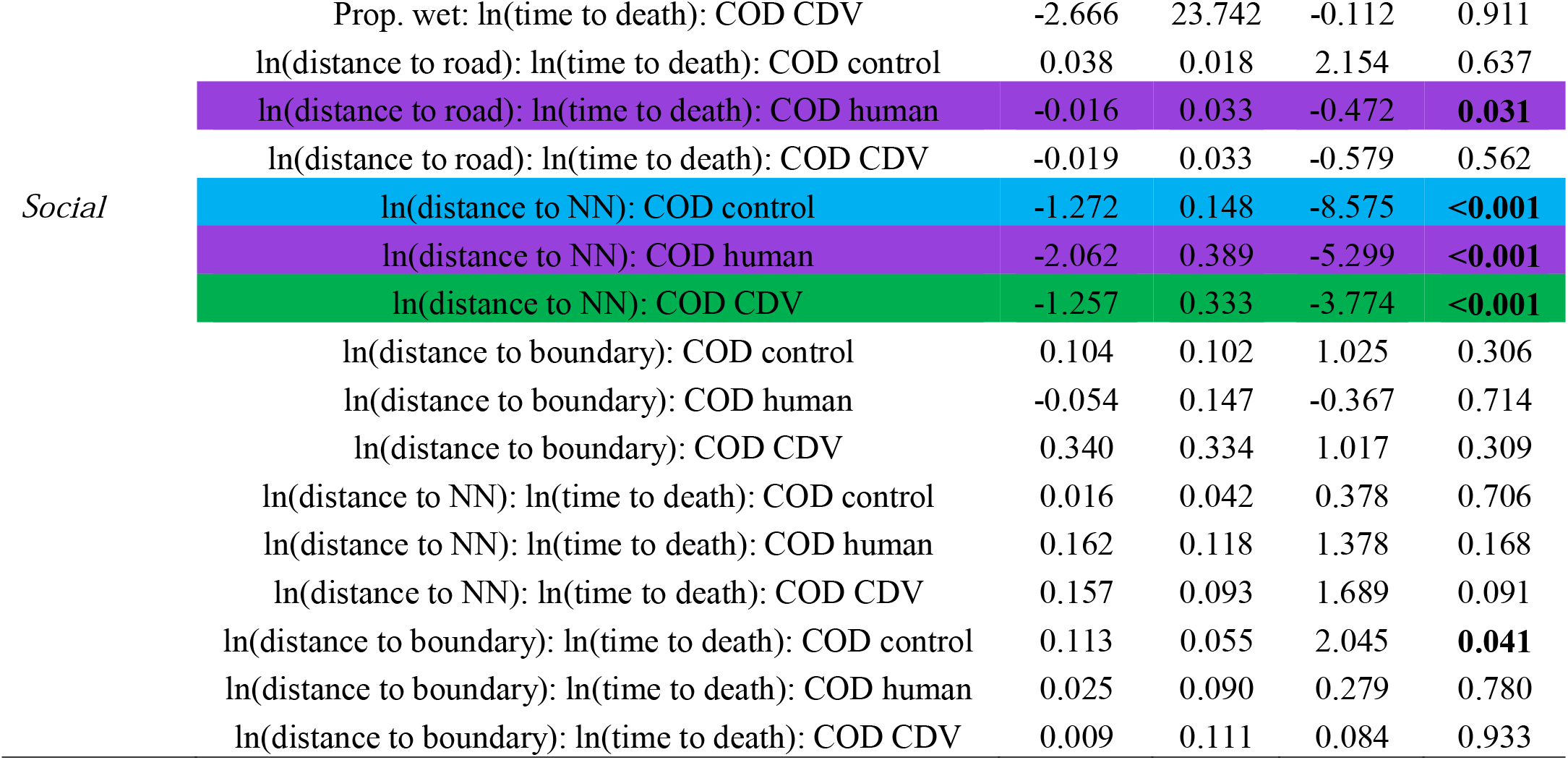
GLMM model output of fixed effects. Bolded p-values are significant at α = 0.05 (n = 29 wolves). Highlighted variables were confirmed to be important with mean speed estimates or relative selection strengths (RSSs) of individuals (blue = control, purple = human, green = CDV).

**Figure 1.**
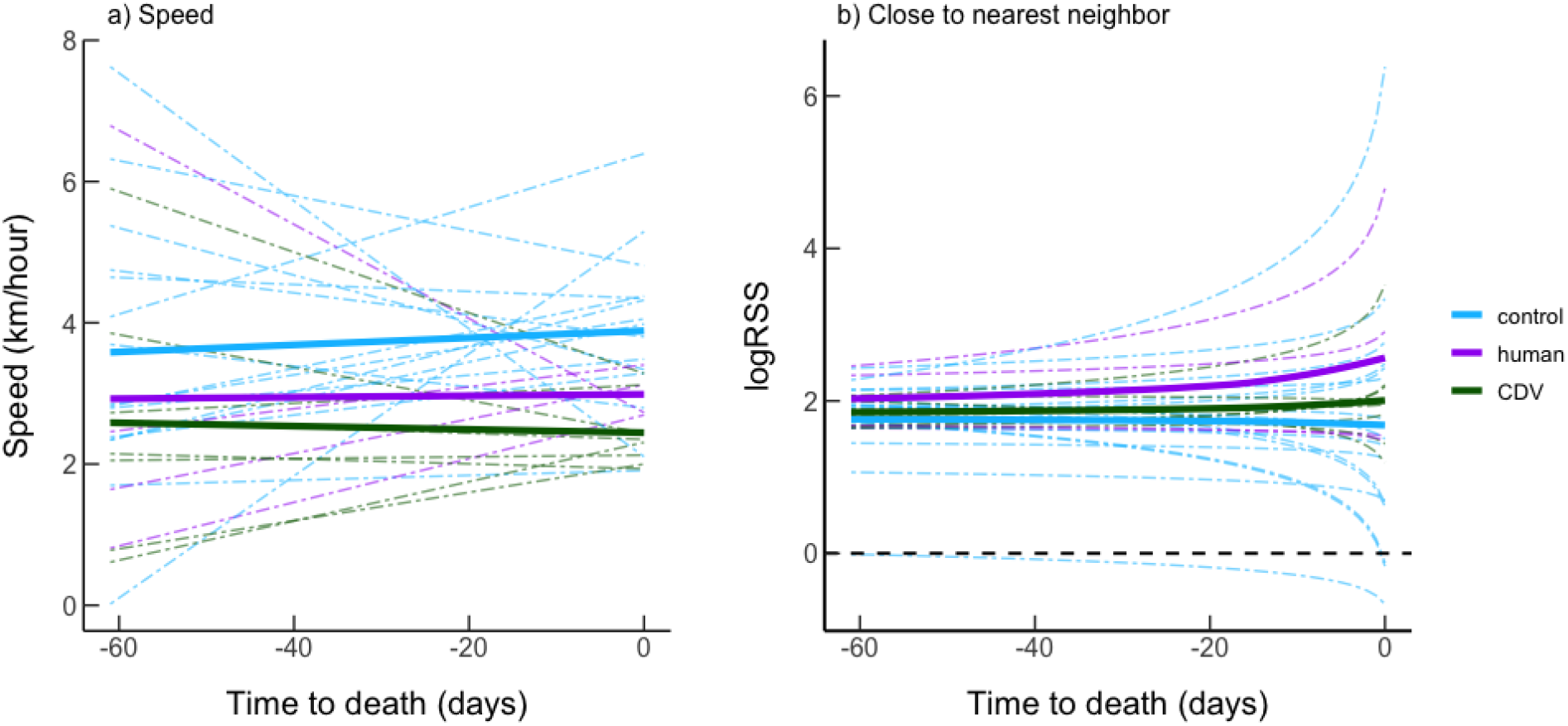
The change in a) speed and b) relative selection strength (RSS) for proximity (250m) to nearest packmate neighbors as wolves that survived (control), died due to humans, and died from CDV neared the end of the two-month period or death. The solid lines represent the population trend with the filled area around the line indicating the 95% confidence intervals, and dashed colored lines represent individual wolves. For the RSS, the black dashed horizontal line indicates no response, above the line represents greater selection than expected, and below the line indicates more avoidance of the habitat type than expected.

**Figure 2.**
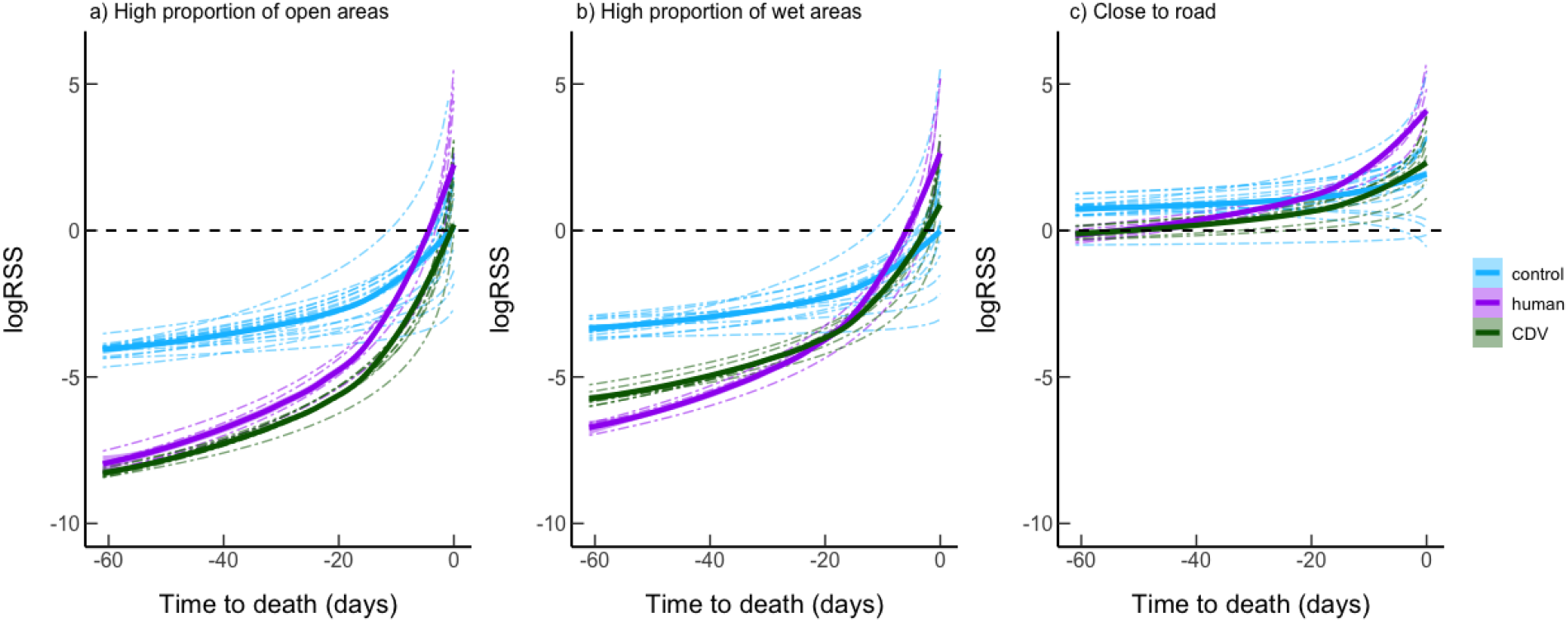
The change in relative selection strength (RSS) for a) high proportion (75% coverage) open areas, b) high proportion wet areas, and c) and proximity (250m) to roads as wolves that survived (control), died due to humans, and died from CDV neared the end of the two-month period or death. The solid lines represent the population trend and dashed colored lines represent individual wolves. The black dashed horizontal line indicates no response, above the line represents greater selection than expected, and below the line indicates more avoidance of the habitat type than expected.

As expected, control wolves did not significantly change their movement patterns, step length and turn angle, over time (Table 1, Fig. 1). Control wolves, however, increased their selection for open habitats as time passed, but showed no change in selection for the proportion of forest or wet habitats as time progressed (Table 1). Control wolves selected to be closer to roads than expected; this did not change over time (Table 1). Control wolves also selected to be closer to their nearest packmate than expected based on availability, and this pattern did not change over time (Table 1, Fig. 1).

In comparison to the baseline behavior of control wolves, human-killed wolves did not show differences in trajectory (cosTA), but they tended to move slower than control wolves (Table 1, Fig. 1). Human-killed wolves did not differ in how they selected for forest, open, or wet habitats compared to control wolves and did not respond to the proximity of roads (Table 1), but they tended to select to be closer to roads right before death (Fig. 2). Human-killed wolves also selected to be closer to their nearest packmate than expected (Fig. 1), but they did not show a significant response to distance from pack boundary (Table 1). However, human-killed wolves spent 21% of their time outside of their territory than in it compared to control wolves that spent approximately 7% of their time outside of their territory (β = 1.61, z = 0.7, p = 0.48). Furthermore, some of the behaviors of the human-killed wolves changed as they neared their eventual death. The human-killed wolves tended to avoid open and wet areas less as they neared death (Table 1, Fig. 2), and they selected to be closer to roads near their death date (Fig. 2). Movement and social covariates did not change as they neared death (Table 1).

Wolves that died from CDV generally moved and changed trajectory more than control wolves (Table 1). The wolves that died from CDV showed no significant differences in baseline response to any habitats compared to control wolves (Table 1). CDV wolves also showed no significant response to road proximity (Table 1). Wolves that died from CDV showed the same behavioral pattern as the human-killed wolves in selecting for be closer to their packmate and showing no significant response to the distance from the territorial boundary (Table 1, Fig. 1). They also spent approximately 5% of their time outside of the territory, not significantly differing from control wolves (β = 1.13, z = −0.6, p = 0.541). CDV wolves also showed changes in movement and habitat selection behavior as they neared their eventual deaths. CDV wolves moved slower and changed direction more the closer they were to dying (Table 1, Fig. 1). Furthermore, CDV wolves tended to avoid open and wet areas less as they neared death (Table 1, Fig. 2). The social behavior of CDV wolves did not change over time (Table 1, Fig. 1).

## Discussion

Our results highlight the interplay among habitat selection, movement, and social behaviors that create and relate to how an individual lives and dies due to human and disease mortality risks. Human-killed wolves did not demonstrate intrinsic, baseline, behaviors but changed their behaviors that increased their risk of death (i.e., selection for roads). Wolves that died from CDV also changed their behaviors as they were dying. Importantly, the increasing selection for open areas and roads and their predilection to spend more time outside of their territorial boundaries compared to control wolves contributed to the risk of death for human-killed wolves. Contrary to our predictions, human-killed wolves were slower than control wolves. Wolves that died from canine distemper virus (CDV) decreased speeds and tended to reduce their avoidance of wet areas as they neared death, as predicted. However, they did not avoid packmates more than control wolves and instead maintained strong selection to be near packmates. Thus, wolves that died from human causes compared to CDV had distinctly different movement and habitat selection patterns. Therefore, the interplay of behaviors provide insight into cause-specific mortality and may help improve our understanding of population dynamics.

Movement and habitat selection behaviors have a synergistic effect so that animals are at greater anthropogenic risk before they die (Lamb et al. 2020). Human-killed wolves selected for habitats where they were at greater risk, consistently moving slower and being more frequently outside of their pack boundary and increasingly selecting for roads before they died. These patterns of selection differed from wolves in our study that were not killed by humans and from general population trends in other predator species. Most notably, carnivores in boreal regions generally select for forested areas and avoid anthropogenic disturbance (Lesmerises et al. 2012; Recio et al. 2019). In many species ranging from predators to prey, variation in consistent individual movement and habitat selection patterns put individuals at greater risk of human caused mortality (M.H. Murray and St Clair 2017; Sawyer et al. 2019). Apex predators move faster in developed areas (Buderman et al. 2018) but tend to slow down and move more cautiously when they hear humans (Suraci et al. 2019). Human-killed wolves may be doing the same as they moved slower than the control wolves and spent time in human-dominated areas near roads. Selecting to be in riskier situations may be a byproduct of life-stage, as the human-killed wolves were all estimated to be 1-2 years old, approximately the age of dispersal (Stronen, Schumaker, et al. 2011). Dispersal is a risky time across species, and the wolves here showed similar movement patterns to other dispersing carnivores (Bartoń et al. 2019; Herrero et al. 2020). Individuals that avoid being killed by humans may learn to better navigate their environment to avoid risky situations (Greenberg and Holekamp 2017). Thus, we see that movement and where individuals select to move synergistically create risky situations that lead to death by humans.

Behavioral responses to disease in particular demonstrated the connections between habitat selection, movement, and social behavior that put individuals at greater risk. Individual wolves that died from CDV contributed to a tragedy of the pack; they maintain their social bonds with their packmates to the detriment of the survival of other individuals in the pack. In our study, most packs died off due to CDV. One pack had three collared wolves die from CDV over the course of a month affecting the pack’s movement for at least two months. Although the risks that arise while infected with CDV may be short-term at the individual level, it has the potential to be more long-term at the pack level where serial infection affects the movements and habitat selection of a pack for months. The social tendency to be close to packmates regardless of illness could influence the inclusive fitness of the group and facilitate continuation of the disease in the population when the whole pack does not disappear. The effects of sickness seem to depend on the type of social interaction. Vampire bats do not change their spatial association behaviors when immune-challenged, but they did decrease their number of grooming partners, likely due to lethargy (Stockmaier et al. 2020 Feb 28). As wolves slow down when they are ill, they likely change their direct social interactions in ways that could not be detected with relocation data. However, the fact that wolves do not change their spatial association patterns indicates that the direct and indirect fitness benefits of social cohesion likely remain greater than the benefits of preventing infecting packmates.

Due to the difficulty of knowing when an animal is going to die and observing it, GPS collar data is useful for quantifying end-of-life behaviors. Behavior-based models have successfully predicted survivorship in population-level models (Stillman et al. 2000), but it is less well understood how individual-level dynamics create those population-level responses. Individual habitat selection, movement, and social behavior are behaviors that can be measured remotely via GPS collars and even camera trapping (Caravaggi et al. 2017). We demonstrate that these behaviors relate to cause-specific mortality. In the future, it would help to pair detailed behavioral observation with movement data to gain more fine-scale insights to disease spread. Furthermore, these remotely measured movement and habitat selection data not only indicate that an animal is dying; they can alert managers to potential risks in the environment such as poisoned carcasses, improper disposal sites, and illegal trapping in our research areas.

The interconnections of habitat selection, movement, and social behavior have individual and population level repercussions (Albery et al. 2020). Here, we relate the interaction of these behaviors to cause-specific mortality after death. Humans harvesting carnivores and CDV outbreaks are both linked to major demographic changes (Cleaveland et al. 2007; Lamb et al. 2020). Thus, fine-scale movement behaviors that are both the cause and consequence of dying are an important factor to consider in conservation and management plans. For instance, scaling up from the individual-level findings here we can get better estimates of the cause of mortality within populations. Furthermore, if management calls for an intervention to control a CDV outbreak, they will have to target members of pack to give a vaccine to stop the spread within the pack once it reaches them (M.J. Silk et al. 2017). Overall, we see that individual-level patterns in the interplay of habitat selection, movement, and social behaviors related to cause-specific mortality from long- and short-term risks.

## Acknowledgements

We would like to acknowledge that our research takes place within the traditional homeland of the Anishinabe people and the Métis Nation, within Treaties 2, 3 and 5 Territory and at the crossroads of Treaties 1 and 4. This work would not be possible without the support of Parks Canada, Manitoba Wildlife and Fisheries Branch, and Manitoba Hydro. In particular we thank K. Leavesley, V. Harriman, T. Sallows, D. Bergeson, K. Kingdon, R. Robinson, and R. Baird for their continuing support. We thank all members of the Wildlife Evolutionary Ecology Lab, including J. Balluffi-Fry, I. Richmond, J. Hogg, J. Kennah, A. Robitaille, J. Aubin, J. Hendrix, Q. Webber, S. Boyle, and L. Newediuk for their reviews. Thanks also to T. Avgar for input on the methods. Thanks to C. Berkvens, who conducted necropsies of wolf specimens. Funding for this study was provided by a NSERC Discovery Grant to EVW.

## Data Availability

Data and code will be made available upon publication on Zenodo and GitHub.

## Competing Interest

The authors declare no conflict of interest

## Author Contributions

JWT, CMP, and EVW conceptualized the idea and study design. CMP, KAK, DLJD, and SZS collected the field data. JWT ran the analyses and drafted the article. All authors critically revised the article and gave final approval for publication.

## Notes

### Competing Interest Statement

The authors have declared no competing interest.

